# High throughput cell mechanotyping of cell response to cytoskeletal modulations using a microfluidic cell deformation system

**DOI:** 10.1101/2024.06.17.599307

**Authors:** Ian M. Smith, Jeanine A. Ursitti, Pranav Majeti, Nikka Givpoor, Megan B. Stemberger, Autumn Hengen, Shohini Banerjee, Joseph Stains, Stuart S. Martin, Christopher Ward, Kimberly M. Stroka

## Abstract

Cellular mechanical properties influence cellular functions across pathological and physiological systems. The observation of these mechanical properties is limited in part by methods with a low throughput of acquisition or with low accessibility. To overcome these limitations, we have designed, developed, validated, and optimized a microfluidic cellular deformation system (MCDS) capable of mechanotyping suspended cells on a population level at a high throughput rate of ∼300 cells pers second. The MCDS provides researchers with a viable method for efficiently quantifying cellular mechanical properties towards defining prognostic implications of mechanical changes in pathology or screening drugs to modulate cytoskeletal integrity.

## INTRODUCTION

Countless cellular functions are regulated by cell mechanical properties, which include a broad spectrum of empirical cellular descriptors and are typically characterized by cellular strain as a response to physical stress. These properties play key roles in a multitude of cellular behaviors across physiological and pathophysiological conditions, from immune cell homing and embryogenesis, to aging and cancer metastasis [1]. Cell mechanotyping, or the quantification of these mechanical properties, provides quantitative measures that can be related to cellular physio-pathology such as degree of differentiation or malignancy [2], [3], [4].

Cellular mechanotyping has evolved over the last 75 years with the continual development of techniques and systems to characterize cellular properties [5], [6], though they generally rely on the same principle: exert a stress on cell, measure their strain response, and use a constitutive equation to determine a “mechanical property” such as an elastic and/or viscous constant. While these techniques have enabled a foundational understanding of cellular mechanical properties, most are technically demanding and low throughput, which has limited their wide-spread use.

For the past 3 decades, the gold standard for quantifying mechanical properties of adherent cells has been atomic force microscopy (AFM) [7], which enables the quantification of the elastic or viscoelastic properties of adherent cells at relatively high spatial resolution [8]. While innovations have dramatically improved acquisition speed, AFM remains a low throughput technique (less than one cell per minute) [7]. Other mechanotyping techniques commonly employed on adherent cells, such as micropipette aspiration or particle tracking microrheology, also have relatively low throughputs with ∼20 to ∼30 cells per hour, respectively, and are further limited by a lack of spatial resolution compared to AFM [6], [9]. Although these methods can provide valuable assessments of cellular mechanics, their low throughput nature and limitation to adherent cells limits their utility in capturing the mechanical heterogeneity of cell populations, especially of those in suspension.

To overcome this problem, microfluidic devices have recently been employed as a high throughput method to quantify population-level cellular mechanical properties of suspended cells [10], [11]. These microfluidic devices incorporate processes commonly used in flow cytometry to apply a hydrodynamic shear stress to aligned single cells [12], [13], [14]; the magnitude of the cellular response determined through image analysis reflects cellular mechanical properties, such as deformability, at a rate upwards of hundreds of cells per second [11]. While the launch of a commercial system supports the promise of microfluidics to mechanotype populations of cells at high throughput, the cost of this commercial option’s instruments and proprietary software are prohibitive [13].

Here we demonstrate a microfluidic cellular deformation system (MCDS) comprised of a commercially available inverted microscope, high speed camera, and programable syringe pump, as well as a custom PDMS deformation chamber. The MCDS was optimized by modifications to the PDMS chamber and fluid viscosity, yielding a microfluidic shear-stress for the efficient population level mechanotyping of commercially available suspended cells at high throughput (∼300 cells/second). We then validated the MCDS by demonstrating its ability to distinguish the impact of established genetic and pharmacologic manipulations reported to alter cytoskeletal properties and cellular mechanics.

## MATERIALS AND METHODS

### Cell culture

MDA-MB-231 (HTB-26), MCF10A (CRL-10317), MCF7 (HTB-22) were purchased from ATCC. MDA-MB-231 cells were grown in Dulbecco’s Modified Eagle’s Medium (DMEM; 11965092, Thermo Fisher Scientific) supplemented with 10% heat inactivated fetal bovine serum (HI-FBS) (10082147, Thermo Fisher Scientific) and 1% penicillin/streptomycin 1000 U/mL (Pen/Strep; 15070063, Thermo Fisher Scientific). MCF7 cells were grown in the same media used for MDA-MB-231 cells, supplemented with 10 μg/mL of insulin (12585014, Thermo Fisher Scientific). MCF10A cells were grown in DMEM/F-12 media (11320033, Thermo Fisher Scientific) supplemented with 5% horse serum (16050130, Thermo Fisher Scientific), 1% Pen/Strep 1000U/mL, 10 µg/mL human Insulin, 100 ng/mL Cholera Toxin (9012-63-9, Sigma-Aldrich), 500 µg/mL hydrocortisone (H0888, Sigma-Aldrich), and 20 ng/mL EGF (SRP3027, Sigma-Aldrich). Cells were washed with phosphate buffered saline (14190144, Thermo Fisher Scientific) and lifted from culture flask using 0.25% Trypsin-EDTA (25200072, Thermo Fisher Scientific). To modulate mechanical properties of cell lines, cells were treated with the phosphatase inhibitor Calyculin A (100 nM for 30 minutes; C5552, Sigma-Aldrich), the Rho-kinase (ROCK) inhibitor Y27632 (20 µM for 30 minutes; Y0503, Sigma-Aldrich), the myosin IIa ATPase inhibitor Blebbistatin (50 µM for 20 minutes; B0560, Sigma-Aldrich), or the microtubule depolymerization inhibitor Taxol (1000 µM for 1 hour; PHL89806, Sigma-Aldrich). Vehicle treatment of DMSO (D2650, Sigma-Aldrich) in its respective concentration was added to cells to function as the technical control.

### MDA-MB-231 viral transduction

We established MDA-MB-231 cells stably overexpressing VASH1 and small vasohibin binding protein (SVBP) to yield an increase in microtubules modified by detyrosination. Control and VASH1/SVBP overexpression vectors were constructed and packaged by VectorBuilder: pLV[Exp]-EGFP:T2A:Puro-EF1A>mCherry (Vector ID: VB010000-9298rtf) and pLV[Exp]-Puro-CBh>mVash1[NM_177354.4]:IRES: mSvbp[NM_001038998.2] (Vector ID: VB211111-1143vqm), respectively. MDA-MB-231 cells were transduced at a multiplicity of infection of ten and supplemented with 10µM polybrene (TR-1003-G, Sigma-Aldrich). 72 hours following infection, the cell culture media was aspirated and replaced with growth media supplemented with the selection antibiotic puromycin at 0.5 µg/mL (A11138-03, Thermo Fisher Scientific). Surviving cells were pooled, expanded, and utilized for downstream experiments as described.

### SolidWorks Flow Simulation

The MCDS AutoCAD sketch file was transferred to SolidWorks to create a three-dimensional device, complete with three holes to represent the MCDS inlets and the outlet. A glass rectangle was connected to the open-faced feature side of the device to represent the irreversible binding to the glass coverslip. The device inlets were then “capped” under the Flow Simulation tab, and the flow rate boundary conditions were added to the defined system. After calculating the resulting flow, pressure and velocity of fluid flowing within the MCDS were visually represented with gradients and streamlines.

### Calculation of theoretical velocity in channel

The theoretical velocity of flow within the deformation region of the MCDS was calculated under the assumption that the system functioned as a Venturi tube. Thus, velocity (v) in the deformation region can be defined as

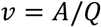

where A is the cross-sectional area of the deformation region of the MCDS, and Q is the cumulative flow rate of the deformation fluid and cell suspension inlets.

### Photolithography

The device mask for the deformation microfluidic device was created in AutoCAD and fabricated by Front Range Photomask. Fabrication of the device negative mold was performed in the University of Maryland Nanocenter Fabrication Lab. To summarize, a layer of SU-8 3010 negative photoresist (Kayaku Advanced Materials) was spin coated onto a silicon wafer (452, University Wafer) to a height of 10 µm. An EVG620 Mask Aligner was used to UV crosslink the SU-8 through the device photomask (Figure 1). Uncrosslinked SU-8 photoresist was washed away using the SU-8 developer (NC9901158, Fisher Scientific). The height of the negative device molds was determined with a profilometer. Devices within 0.25 µm were used in testing. Finally, wafers were silanized following plasma treatment (PDC-001, Harrick Plasma) by incubation with tridecafluoro-1,1,2,2, tetrahydrooctyl-1-trichlorosilane (78560-45-9, Gelest) overnight in a vacuum desiccator.

**Figure 1:**
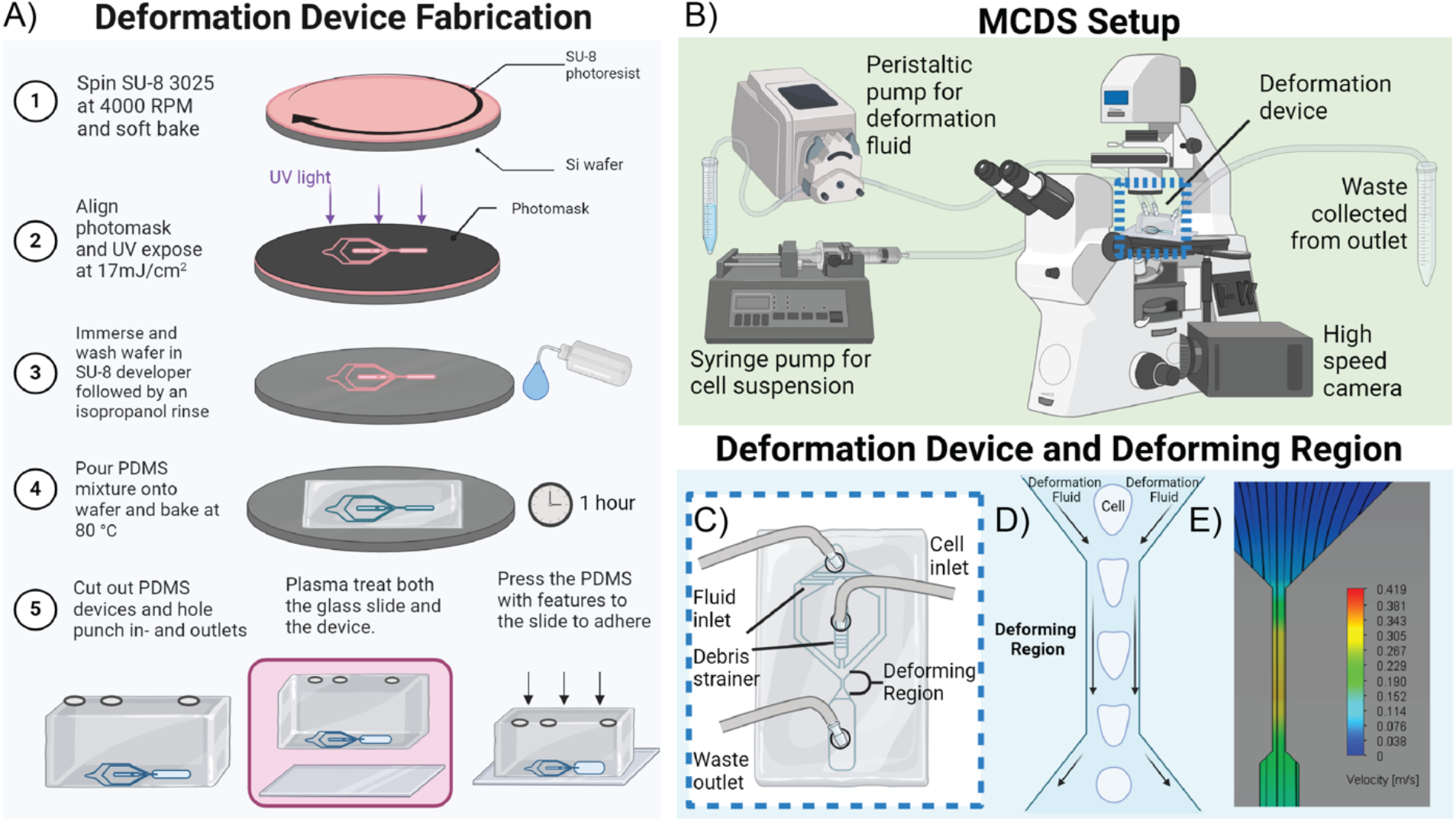
Graphical overview of MCDS fabrication, setup, and mechanism of action. A) representation of the methods used for the fabrication of the MCDS positive mold, the PDMS casting of the mold, and the irreversible bonding of the device to a glass coverslip. B) Graphical representation of the system setup. C) An up-close look at the box highlighted in B, detailing the device components and tubing setup. D) a graphical representation of the cell morphology at different distances within the deformation region of the device. E) A SolidWorks FlowView representation of flow velocity and streamlines within the MCDS.

### PDMS and glass adhesion

Polydimethylsiloxane (PDMS) was mixed at a 10:1 base polymer to crosslinker (1317318, Dow). This mix was poured over the silicon wafer, which was placed in a vacuum desiccator for 30 minutes and then baked at 80^°^C for at least 1 hour. The PDMS was removed from the wafer, cleaned with ethanol and water, dried at 80^°^C for five minutes, and then plasma treated for two and a half minutes. The PDMS devices were then bonded to glass coverslips (which were also cleaned, and plasma treated simultaneously with the PDMS) under pressure overnight at 80 ^°^C. The following day, devices were plasma treated again for two and a half minutes and 10 µL of 8% Pluronic F68 (P1300, Sigma-Aldrich) was added to the device to prevent cell sticking. Devices were then washed three times with PBS. For long term storage, the devices were soaked in DI water, put in a closed container, UV treated, and left in the fridge for up to three months.

### Preparing cells for the MCDS

Following detachment from culture flasks, cells were strained (76327-098, VWR), centrifuged, and resuspended to count. Cells were centrifuged again, and the resulting pellet was resuspended with 10 uL of PBS. Deformation fluid was then added to the suspension to produce a concentration of 4×10^6^ cells/mL. Deformation fluid consisted of 0.1 wt.% Sodium Azide (S2002, Sigma-Aldrich), 0.5 wt.% BSA (A4612, Sigma-Aldrich), and 0.5 wt.% methylcellulose (M7027, Sigma-Aldrich) in a PBS. This cell suspension was transferred into a 3mL syringe (53548-017, VWR), which was then attached to the cell inlet tubing for the device system. The cell suspension was pushed into the tubing, and the 3 mL syringe was loaded with fresh deformation fluid to increase the cell density going through the device.

### Preparing the MCDS

The deformation system (Figure 1B) consists of an Olympus IX71 microscope configured with a high-speed Phantom MIRO C210 High Speed Camera. During acquisition, the camera imaged at 2,000 frames per second (fps) with an exposure time of 7 µs as optimized. Microscopy was done using phase contrast with a 20x objective lens. The deformation fluid (see *Preparing cells for the device*) for inducing the hydrodynamic shear stress was perfused via a peristaltic pump at a rate of 300 µL/hour. The cell suspension was prepared as described in *Preparing cells for the device* and perfused through the system via a syringe pump at a rate of 300 µL/hour. The volumetric flow rates at which fluids were perfused was determined via MCDS optimization to output observable cell deformation without compromising cell health. Following initiation of flow into the device, there was a five-minute wait step to allow the system to equilibrate. The cellular deforming region of the system was then imaged for five seconds. Image sets were exported as Tiff files.

### Quantification of deformation fluid viscosity

Four stock solutions of deformation fluid were created with different concentrations of methylcellulose (0 wt.%, 0.25 wt.%, 0.5 wt.% and 1 wt.%). These solutions were filtered (0.2 µm PES, 430771, Corning) and 1 mL of each was warmed to room temperature. The solution was then perfused through the RheoSense microVISC Handheld Viscometer to calculate sample viscosity.

### Western blotting

Equal numbers of MDA-MB-231 cells were plated in 6-well dishes (3×10^5^ cells per well) and processed for Western analysis one day following plating. Briefly, cultures were washed twice with PBS and each well was collected into an equal volume of 2x Laemmli sample buffer (1610737, BioRad), sonicated and heated before being processed via SDS-PAGE (,4-20% Mini-PROTEAN® TGX™ precast gels, 4561094, BioRad). Proteins were transferred to a membrane (Immobilon-FL PVDF, IPFL00010, Millipore), stained with Revert 700 Total Protein Stain (926-11011, LI-COR Biotech) for five minutes at room temperature, rinsed in ultrapure water and imaged on the LI-COR Odyssey CLx system. Immediately after imaging, the membrane was rinsed in ultrapure water and blocked (SuperBlock PBS 37515, Thermo Fisher Scientific) for one hour at room temperature. Membranes were probed overnight for β-tubulin (T4026, Sigma-Aldrich), detyrosinated-tubulin (31-1335-00, RevMAb Biosciences) and tyrosinated-tubulin (1864-1, Sigma-Aldrich). Blots were washed three times for five minutes with 1x TBS + 0.1% Tween 20 (TBST), incubated with the corresponding secondary antibody (1:7500 in TBST) for 1 hour at room temperature, washed three times for five minutes with TBST, and imaged on the LI-COR Odyssey CL-x system.

### Data analysis

Cells were manually traced and analyzed using FIJI (ImageJ). After determining cellular borders, cell shape factors including area, perimeter, circularity, and major and minor axis lengths were quantified. At least 100 cells were analyzed from each video file. Cells with the following traits were excluded from analysis: cells dragged along the cell wall, cells with noticeable blebbing, cells in contact with other cells, multinucleated cells, and cells with distinct blebbing (Supplemental Figure 1).

### Statistical analysis

Unless otherwise stated, any graph comparing more than two groups was analyzed in Prism GraphPad with the non-parametric Kruskal-Wallis test followed by Dunn’s multiple comparisons test. Graphs with two groups were analyzed by a Mann Whitney test (* p<0.05, ** p<0.01, *** p<0.001, **** p<0.0001). Experiments were conducted in triplicate and resulting values were pooled between trials. Unless otherwise shown, these pooled trials were displayed in violin plots to visually present the distribution of the large numeric data sets, with the center line representing the mean.

## RESULTS

### Microfluidic Cellular Deformation System was developed and modeled

The MCDS chamber design, innovated upon one previously published [13], was fabricated via conventional processes of photolithography, PDMS casting, and irreversible bonding to glass cover slips through plasma treatment (Figure 1A). Following traditional methods for cell proliferation and preparation for flow cytometry, the cell suspension and deformation fluid were added to their respective tubing and connected to the MCDS. Following initiation of flow and a wait step for the system to equilibrate, cell deformation was imaged with the Phantom high-speed camera (Figure 1B-D and Supplemental Video 1). Models of the flow velocity (Figure 1E) and pressure (Supplemental Figure 2) within the deformation region of the MCDS were created with SolidWorks FlowSimulator.

### MCDS validation revealed its robust ability to deform cells

In initial trials of the device, MDA-MB-231 cells were used since their mechanical properties (both in suspended and adherent states) have been previously characterized [15], [16], [17], [18]. Cells flowed through the device visually displayed deformation when compared to suspended cells (Figure 2 A & B). Upon entering the deformation region of the MCDS, cells experienced the greatest deformation in the first third of the channel, followed by a gradual decrease in deformability in the latter two thirds of the channel (Figure 2B). This deformation was in line with the gradually reduced pressure differential within the channel (Supplemental Figure 2). Three metrics were used to quantify change in cell shape within the device: (1) deformation, (2) deformation versus cell area, and (3) the ratio of fit ellipse major axis over minor axis. There was a significant increase in the deformation for cells in the MCDS in comparison with suspended cells in static conditions (Figure 2C-E).

**Figure 2:**
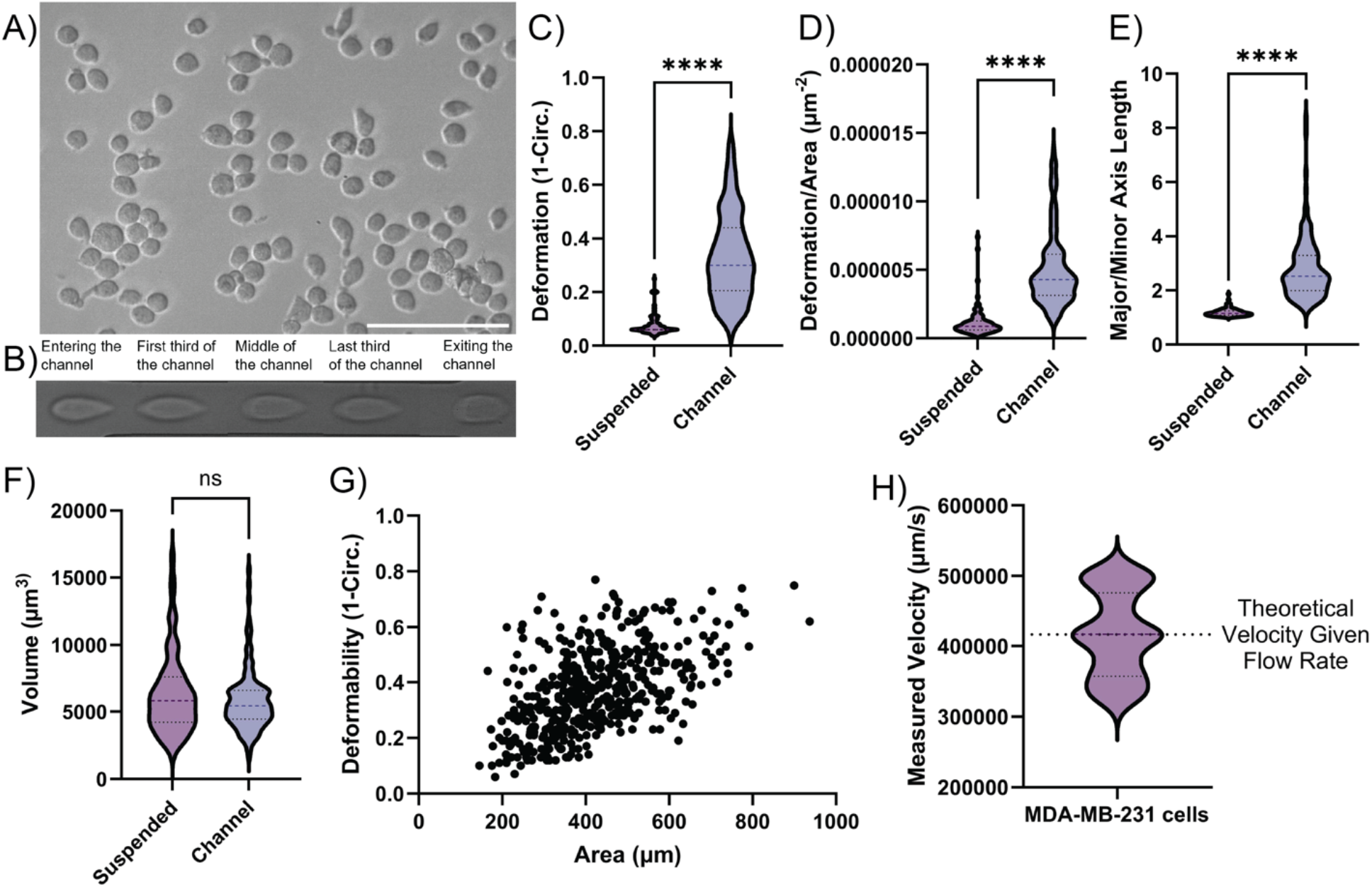
Cellular deformation of MDA-MB-231 cells is detectable in MCDS: A) MDA-MB-231 cells suspended in deformation fluid images on the MCDS microscope. B) A single representative image of a MDA-MB-231 cell being flowed through the channel, where positions of the cell within the channel are stitched together to show deformation at each region within the channel. C-E) Comparison of suspended and deformed cells using quantifiable metrics previously used in deformability cytometry (n = 125). F) Calculated volume for cells within suspension and cells in the deforming region of the MCDS; assuming spheres and ellipsoidal bodies (n = 125). G) Dot plot showing the resulting deformation given the area of MDA-MB-231 cells within the MCDS (n = 500). H) Velocity of MDA-MB-231 cells traveling through the device, calculated by hand, compared to the theoretical velocity calculated from the pump’s flow rates.

The reproducibility of cell tracing was identified by analysis of the same videos across a single treatment by four of the authors (Supplemental Figure 3). As there was a subtle variability in the analysis between the authors, all subsequent data presented here were analyzed by a single person. To ensure an unbiased analysis, all files were blinded for analysis; however, following acquisition of data from the blinded data, a randomly selected group was unblinded and reanalyzed with the bias of knowing the treatment, and thus expected deformation of cells. From this test, there was no significant difference in analyzed deformation between blinded and unblinded groups (Supplemental Figure 3).

To further validate the device, theoretical results of cell shape descriptors and flow characteristics were compared with the results collected within the MCDS. First, the volumes of the cells calculated based on images of cells in suspension or under deformation within the MCDS (assuming spherical and ellipsoidal shapes, respectively) were not statistically different (Figure 2F). No significant change in volume suggests that the cells were not ruptured or damaged upon entering the deformation region of the MCDS. Furthermore, a positive trend of cellular deformation versus cell area was shown (Figure 2G). Finally, the velocities of cells moving within the channel were compared to the theoretical velocity (assuming the system to be a Venturi tube) (Figure 2H). The average velocity of the cells determined from the MCDS was almost identical to the theoretical velocity. Additional cellular parameters within the deforming region, such as cell distance from edge, axis lengths, and aspect ratios were also calculated (Supplemental Figure 4). These shape descriptors provide further evidence of the deformation of cells within the MCDS when compared to those in suspension.

### Optimizing the MCDS

In addition to the need for the MCDS to induce cell deformation, other engineering requirements included 1) limiting cell death or stretching, 2) inducing a consistent distribution of measurable cellular deformations, 3) producing stable flow, and 4) creating reproducible conditions within the device to allow for consistent analysis. Previous literature on deformability cytometry specifications informed our system parameters used in initial prototyping stages, including flow rates, viscosities, system equilibrium times, etc. [13]. However, when these previously defined system parameters were applied to our MCDS, there were problems with cell health, acquisition of videos, and reliable collection of data; thus, optimizing these, and other MCDS parameters associated with image acquisition, was essential for the collection of viable and reproducible data.

In some instances, the cells traveling through the deforming region of the MCDS underwent rupturing or excessive cell stretching (Supplemental Figure 1), likely due to the high hydrodynamic shear stress of the deformation fluid. To ensure the device met the engineering requirements of maintaining cell health, while inducing quantifiable cellular deformations, the flow rates into the MCDS were refined. To optimize these flow rates, cells in a standard deformation fluid were flowed through the MCDS with a five-minute wait step after initiation of flow to allow the system to equilibrate before data collection. Three flow rates of the cell suspension were assessed in combination with three flow rates of the deformation fluid, and the resulting cell deformations were evaluated (Figure 3A). There was an upward trend in deformation as the flow rates of both the cell suspension and the deformation fluid increased.

**Figure 3:**
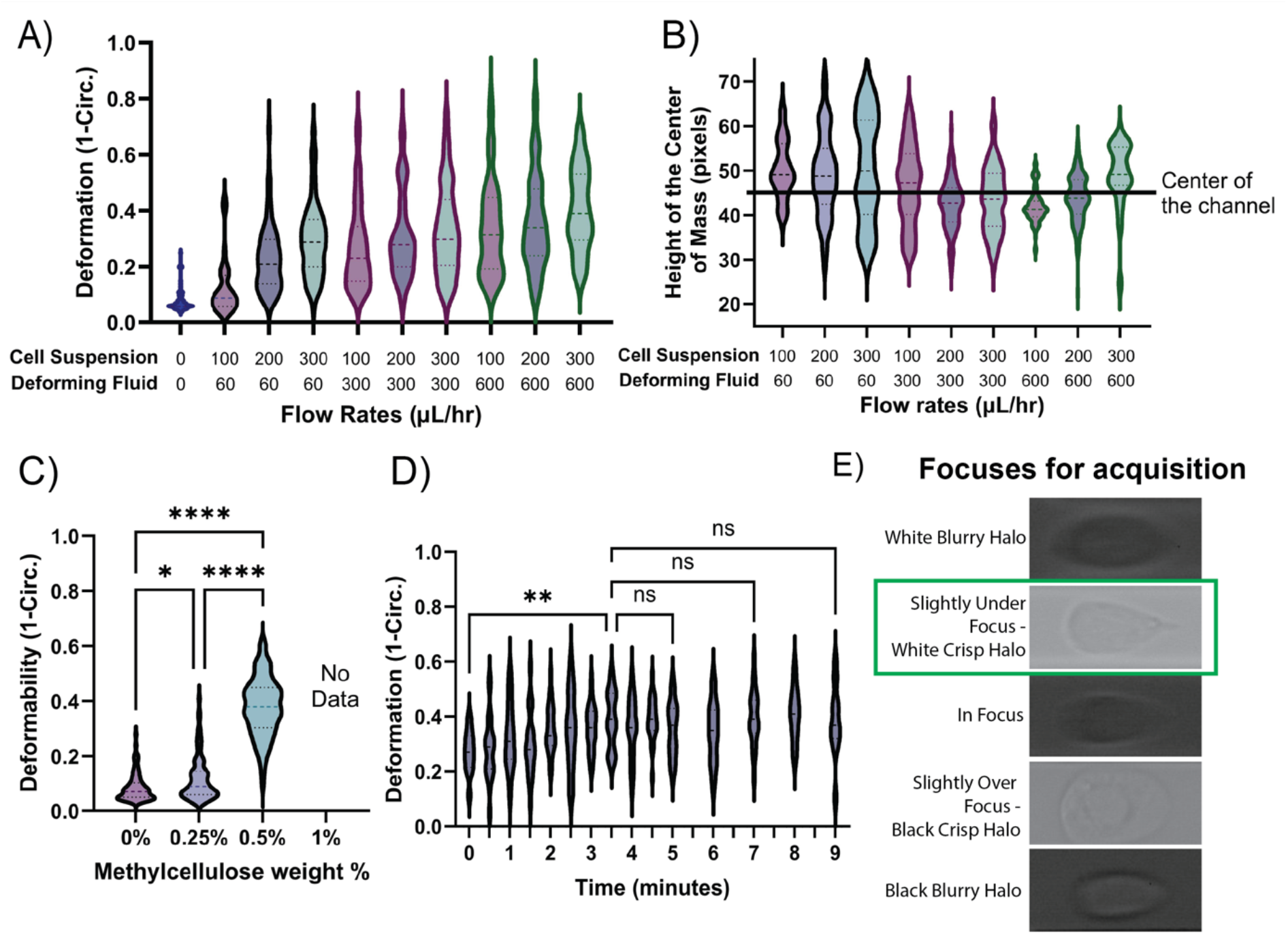
Optimizing MCDS parameters to achieve viable sample collection for efficient analysis. A) The deformation of MDA-MB-231 cells given the flow rates in µL/hour of the cellular and deformation fluid inlets; these are represented as the first & second numbers for each group on the x-axis (n= 125). B) Location of the center of mass of analyzed MDA-MB-231 cells within the deforming region given flow rates in µL/hour (n = 125). C) Deformation of MDA-MB-231 cells in deformation fluids of varied viscosities created by adding methylcellulose at different weight percentages. The 1% methylcellulose column is labeled with no data, as cells entirely ruptured upon entering the channel, thus no data could be collected (n = 125). D) Deformation of MDA-MB-231 cells given time after the initiation of cell and deformation flow (n = 25). E) Representative images of cells in focus within the deforming region of the MCDS to ensure consistent data acquisition for reproducible analysis. The green box represents the foci chosen for future experiments; however, the black crips halo would also be a viable choice for consistent analysis.

The initial flow rates used in the prototyping stage of the device yielded excessive cell death. Furthermore, cells were often pushed up or down against a wall of the deformation device, likely due to the unstable flow. A previous study noted unstable flow in the constricting region of a similar microfluidic device following at high flow rates, resulting in non-streamline cell movement [19]. Therefore, in addition to having flow rates that induce viable and effective cell deformation, our MCDS needed to meet the engineering requirement of achieving stable flow rates that center cells in the channel. This streamlined movement in the center of the channel needed to achieve symmetrical cellular deformation which would further enhance reproducibility of the MCDS. Quantifying the distribution of the center of mass of cells referenced to the center of the channel revealed variation amongst the different flow rates (Figure 3B). From these results, the flow rate of 300 µL/hour for both the cell suspension pump and deformation fluid pump was chosen because they induced a Gaussian distribution of deformation data points, displayed little cell rupturing, and resulted in a center of mass average closest to the center of the channel.

To further prevent cell rupturing or stretching under shear stress, the effect of deformation fluid viscosity on cell deformability was determined within the MCDS. Deformation device fluid stocks were made with 0 wt.%, 0.25 wt.%, 0.5 wt.% and 1 wt.% methylcellulose, resulting in solutions of varying viscosity (Supplemental Figure 5), with the standard deformation fluid being 0.5 wt.%. They were run through the MCDS with the flow rates of 300 µL/hour with videos captured five minutes after the system reached equilibrium. When the deformation fluids of different viscosities were used in the device, there was a significant increase in cell deformation between 0 wt.% to 0.5 wt.% methylcellulose (Figure 3C); however, with the higher viscosity deformation fluid of 1 wt.% methylcellulose, the shear stress experienced by cells in the deforming region was excessive to the point of causing all cells to rupture. Since 0.5 wt.% methylcellulose induced a large distribution of deformation without causing aberrant cell health, it was chosen as the viscosity value for the remaining experiments.

From there, the next engineering requirement for the MCDS was to ensure that its setup and resulting data for processing were reproducible. One factor of reproducibility was the time for the system to equilibrate and reach a steady state flow to induce consistent deformation values. To accomplish this, deformation videos were captured every subsequent 30 seconds for the first five minutes; videos were then captured every minute between five and nine minutes after initiation of flows (Figure 3D). The five-minute mark was chosen as the switch point for sampling rates as it was shown to be a time where previous deformability cytometry systems reached an equilibrium [13]. From 0 to 3.5 minutes, there was a significant increase in the deformation of cells, which leveled off, showing no further significant changes in deformation for latter time points. Considering this data, the time for MCDS equilibrium before acquisition of videos was optimized from five minutes to four minutes in subsequent cell treatment experiments.

Additionally, MCDS reproducibility was ensured by producing consistent output videos to control variation during analysis. A potentially large point for variation during acquisition of deformation videos is the focus of cells within the MCDS, as this could lead to discrepancies in the visual definition of cell borders [11], [12], [13]. Of the five different focal planes of cells observed within the MCDS, two provided the analyzer with clearly defined borders to accurately trace the cell (Figure 3E). These borders were caused by the cells being slightly over- or under-focused, resulting in black and white halos, respectively, surrounding the cells. Though either could be chosen for analysis, to ensure consistency, all cells in further experiments were focused at a slightly under-focused level, resulting in the clear white halo around the cell border (Figure 3E). Further parameters such as cell density, cell staining, image brightness, or acquisition rate were evaluated for device optimization, but these had no relevant impact on any of the engineering requirements (Supplemental Figure 6).

### Validating the MCDS with mechano-modulating drug treatments and defined cell types

Following development and optimization, we validated the ability of the MCDS to differentiate altered cell mechanics in response to established pharmacology targeting either microtubules or the actomyosin cytoskeletal components [20], [21], [22]. The concentrations of these drugs were chosen as they have significant changes on MDA-MB-231 mechanical properties in adherent and suspended states. Blebbistatin and Y27632 inhibit myosin IIa activity, albeit with differing mechanisms of action. Consistent with reports that reduced myosin IIa activity in suspended cells yields increased cell mechanics due to reduce cortical actin turnover [23], the MCDS found both drugs reduced cell deformation. Aligned with the myosin IIa-actin axis we also show that Calyculin-A, a broad-spectrum phosphatase inhibitor that increases in myosin IIa activation, increases cell deformation.

The MCDS was further validated by treatment of MDA-MB-231 cells with Taxol, a drug that prevents microtubule depolymerization[20] yielding the robust proliferation of microtubules that are dramatically enriched in tubulin post-translational modification’s (PTM) (i.e., detyrosination and acetylation), changes that together yield increased cell mechanics[24]. Consistent with these reports we show that MCDS differentiated a measurable reduction in the deformability in Taxol-treated cells vs control. Together, these results validate the MCDS as a system capable of delineating the effects of cytoskeletal modifications on cellular deformability.

In addition to pharmacologically modulated cell mechanical properties, cellular deformation was expected to change as a function of cell type. It is well established that as cells move towards a more metastatic state, their stiffness decreases and their deformability increases [2], [18], [25], which aids in their invasion throughout the metastatic cascade. This biophysical phenomenon has been observed in the commonly used trio of breast epithelial cell types, including non-tumorigenic MCF10As, non-metastatic MCF7s, and highly metastatic MDA-MB-231 cells. These cell types have commonly been studied together to discern mechanical properties as a function of tumorigenicity; hence, we chose them for validation of the MCDS [26]. As expected, there was a significant increase in deformability of the highly metastatic MDA-MB-231 cells in the MCDS when compared to their less metastatic counterparts (Figure 4C-D). There was no detectable change in the deformability between MCF10A and MCF7 cells.

**Figure 4:**
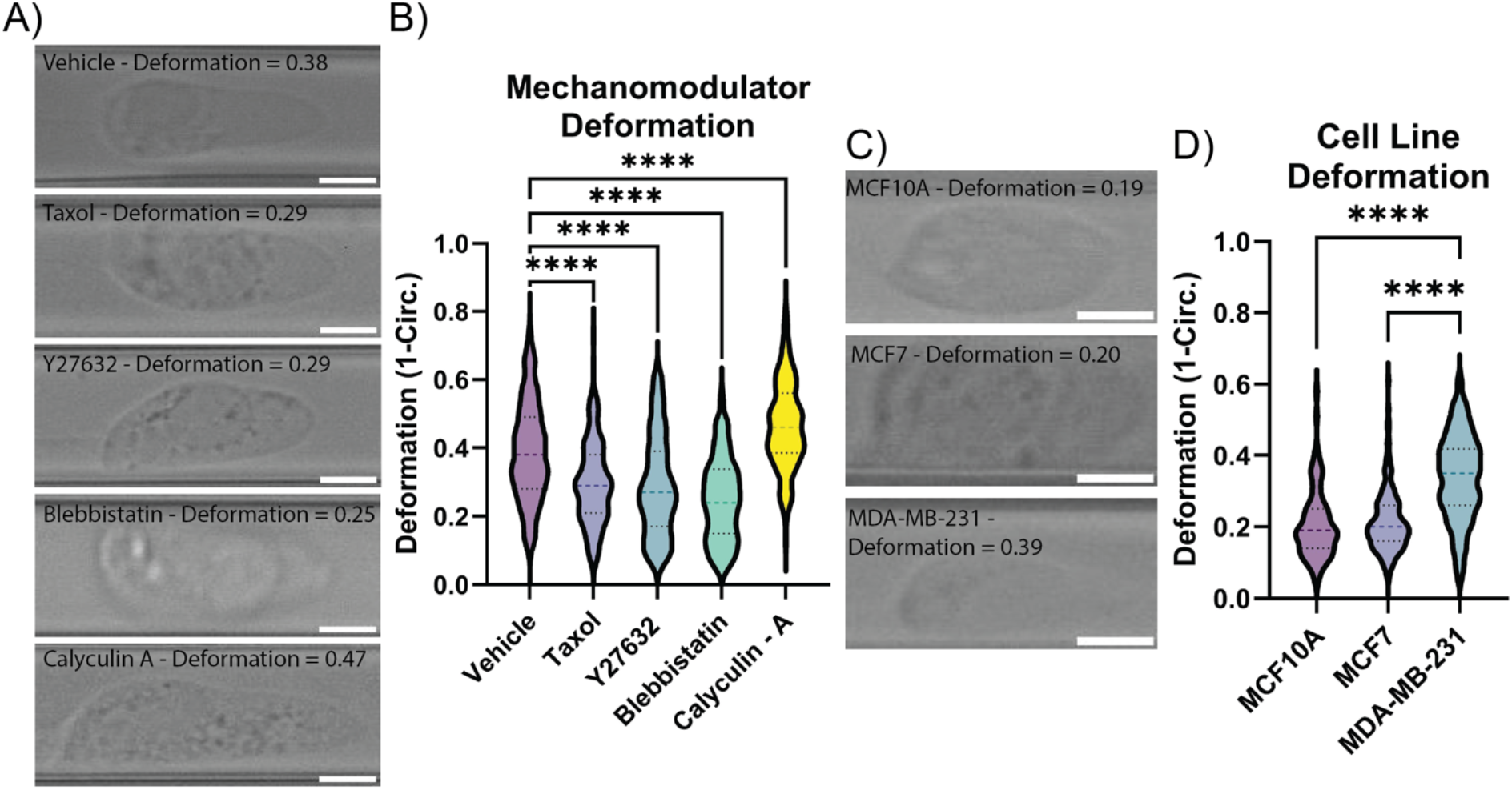
Validation of MCDS with predefined treatments for stiffness: A) Deformability of MDA-MB-231 cells given treatment with Vehicle, Taxol (1µM), Y27632 (20µM), Blebbistatin (50 µM), and Calyculin-A (100nM) (n=200). C) Deformability of non-tumorigenic MCF10A, non-metastatic MCF7, and highly metastatic MDA-MB-231 cells. (n=125). B & D) Representative images from each treatment or cell line with a deformability at approximately the average deformability for that cell type. Scale bar = 10 µm. ****p<0.0001

### Validating the MCDS in cells genetically overexpressing VASH1

The structure and properties of microtubules are regulated by PTMs to their tubulin proteins. Detyrosination (deTyr) is the reversible cleavage of the α-tubulin C-terminal tyrosine (Tyr) residue by VASH1/SVBP or VASH2/SVBP [27] or the more recently identified MATCAP [28]. This PTM, termed deTyr-tub, promotes the microtubule association with multiple protein binding partners which increases cytoskeletal connectivity and thus cell mechanics [29].

Our group has made discoveries on deTyr-tub as a regulator of cellular physiology and pathology in cancer and striated muscle disease. Our works in mature striated muscle cells (skeletal muscle and heart) established deTyr-tub as a regulator of cytoskeletal mechanics and implicated the disease dependent increase in deTyr-tub for the deleterious increase in cell mechanics that drives pathology[30]. Further work used the genetic overexpression of VASH 2 + SVBP in WT muscle to validate the independent impact of deTyr-tub to these effects. In that work, measures of mechanical properties were collected via AFM to quantify the alteration in cell mechanics. Here the low throughput nature of this technique limited the number of cells that could be efficiently analyzed. Unfortunately, the large size and asymmetric shape of these primary muscle cells make it impossible to mechanotype them within the MCDS.

Prior to our work in striated muscle, our group discovered elevated deTyr-tub as a pathologic adaptation in cancer cells that is found at invasive tumor fronts [31]. Motivated by our evidence linking elevated deTyr-tub to increased metastatic potential [31], [32], [33], and new evidence of increased cell mechanics by MCDS (Figure 4C & D), we sought the independent impact of deTyr-tub to these increased cell mechanics.

To this end, we developed a stable MDA-MB-231 cell line stably over expressing VASH1 - SVBP (VASH1 - OE). Consistent with this enzyme’s carboxypeptidase activity promoting detyrosination, western blotting showed a significant increase in deTyr-tub, a concomitant reduction in tyrosinated tubulin (Tyr-tub), and no change in tubulin expression in the VASH1 – OE line compared to the MDA-MB-231 GFP control cell line (Figure 5A). Consistent with deTyr-tub as a positive regulator of cytoskeletal mechanics, we show a significant reduction in MCDS deformation of the VASH1 - OE verses the control cell line (Figure 5B).

**Figure 5:**
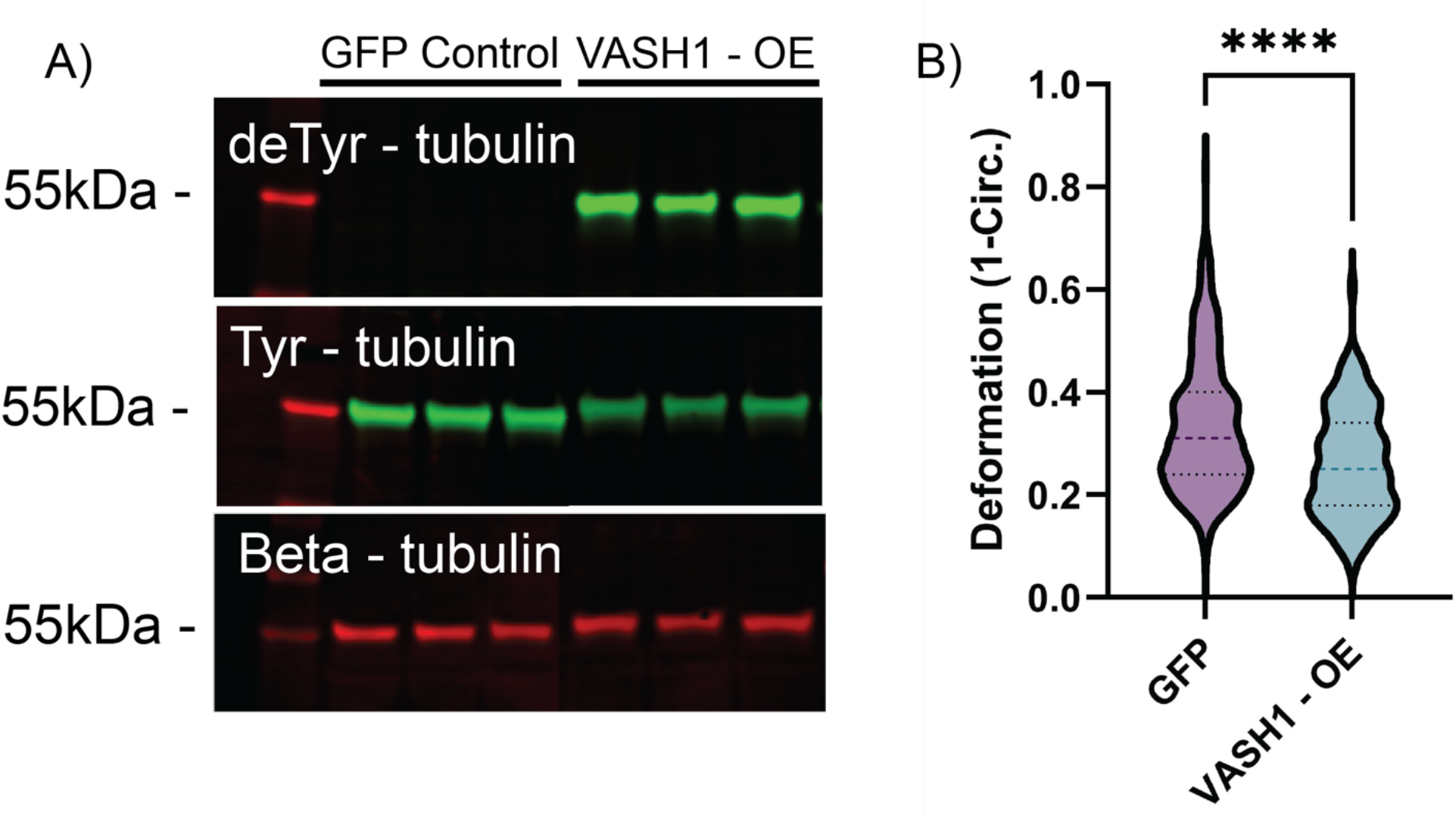
Detyrosinated microtubules reduce cell deformability. A) western blot of deTyr-, Tyr-, and β-tubulin for control and VASH1-overexpressing cells. B) Deformation of MDA-MB-231 control and VASH1-overexpressing cells (1-way ANOVA, ****: p<0.0001).

## DISCUSSION

Quantification of cellular mechanical properties has become a foundational for defining cell behaviors and phenotypes [6], with properties like cell deformation that can be predictive of physiological and pathological states [3], [4]. While commercial instruments are available to quantify cellular mechanical properties in populations of suspended cells, the high cost, extensive trainings, and proprietary nature [11], [13] can be significant barriers to adopting these methods. We have overcome these challenges with the design and validation of the MCDS (microfluidic cellular deformation system) for the efficient quantification of single cell mechanical properties at a population level.

Our first design goal was creating an MCDS comprised of components largely available in most research groups. The optical platform was comprised of a repurposed inverted microscope coupled to a new high-speed brightfield camera capable of ultra-high frame rates. Our fluidics system was reproposed programable syringe pump, and our deformation chamber was custom fabricated from PDMS to optimize the deformation of MDA-MB-231 cells, a line whose mechanical properties have been extensively quantified and characterized with numerous methods in both adherent and suspended states.

Our initial metric for design success was observable and quantifiable cell deformation when compared to suspended cells (Figure 2 A-E). Resulting shape metrics of cells within the MCDS were compared to theoretical phenomena, and all data supported the initial hypotheses: cells did not rupture or lose volume in the channel (Figure 2F), cell deformation increased in relation to increasing shear stress (Figure 2G), and the average velocity of cells within the MCDS very closely resembled the theoretical velocity calculated by hand (Figure 2H). Though no publications have employed deformation cytometry to mechanotype MDA-MD-231s specifically, previous work has compared shape descriptors of other cells in deforming devices to their suspended counterparts. These authors report a significant increase in the cell deformability metric, which is quantitatively similar to our current results [13]. The MCDS however, realized a larger magnitude and distributions of cell deformability than this previous study, which may have arisen from technical design differences, or the larger cell type used in this study.

To ensure our system has viable, reproducible, and rigorous results, engineering parameters for the MCDS were set. While initial trials were conducted with parameters reported in literature, [13] they were optimized to maximize the reproducibility of data acquisition and analysis. To ensure our system has viable, reproducible, and rigorous results, engineering requirements were set, and variable system parameters were optimized.

Optimizing the flow rates of the deformation fluid and cell suspension addressed three of the engineering requirements set for the MCDS: creating a system that 1) produces a stable flow profile with a reproducible of cell position through the flow, 2) induces a consistent distribution of measurable cellular deformation, and 3) elicits minimal cell damage or death. Quantifying cell deformation while iteratively altering MCDS inlet flow rates, we identified 300 µL/hour for both inlets as a parameter that reliably induced cell deformation in a standard distribution, while maintaining cell viability (Figure 3 A). To further ensure uniform and streamlined shear stress in the MCDS for symmetrical cellular deformation, the average height of the center of mass of cells was calculated and compared to the ideal center of the deforming channel (Figure 3B). This effect directly impacts the reproducibility of results derived from deformation cytometry systems and should be considered across all systems for future work in the field.

The engineering requirement for cell viability within the MCDS was also investigated by modulating the viscosity of the deformation fluid used within the system. Increasing the viscosity resulted in increased shear stress on the cells, until the point of cellular rupture (Figure 3C). The concentration of 0.5 wt.% methylcellulose was chosen at the defined flow rate as it allowed for the observable deformation, which could be captured within our system. However, the rationale for testing this effect is cyclical, as the flow rates determined from the previous step were optimized with the 0.5 wt.% methylcellulose stock, and thus would likely be the best condition in this test group. For this study, the above conditions satisfied the specified engineering requirements.

The MCDS also had to meet the engineering requirement of a system that 4) creates reproducible conditions within the device to allow for consistent analysis. Upon optimization, it was found that the MCDS reached an equilibrium with non-significant differences in cell deformability around a minute and thirty seconds faster than previously noted, thereby allowing for a shorter wait step before the acquisition of data (Figure 3D). Considering previous findings, the microscope focus of cells within the MCDS impacts the analysis process (Figure 3E); thus, all cells were captured under a consistent and defined focus to reduce analysis variability.

Having established the reproducibility of the MCDS, we next rigorously tested the ability of the MCDS to reliably quantify cell mechanics in populations of non-adherent cells across established cell types and pharmacologic and genetic strategies reported to alter cell mechanics. First, we showed that the MCDS reliably differentiated the mechanical properties of MDA-MB-231 cells challenged with actin or microtubule targeted pharmacological approaches shown to either stiffen of soften the cytoskeleton (Figure 4A & B). Importantly, these results were qualitatively similar to those reported with similar pharmacology [23].

We next sought to confirm the inverse relationship between the metastatic potential of cancer cells and their cell mechanics [16], [25], [34], which enhances invasion by enabling cells to squeeze and migrate through their surrounding tissue [35]. As previously noted, the mechanical properties of highly metastatic MDA-MB-231 cells have been widely characterized across adherent (AFM, Micromanipulator, magnetic bead twisting) and suspended (optical tweezers, deformation cytometry) states [15], [16], [25], [36], [37]. These MDA-MB-231 cells are reported as being softer than both the non-tumorigenic MCF10A or the non-metastatic MCF7 cells [26], which have been consistently identified as having little to no difference in stiffness between each other [15], [38]. Our ability to distinguish these differences in cell mechanics across these cell types further confirms the efficacy of the MCDS (Figure 4C & D).

Lastly, we determined the cell mechanical consequence of a genetic increase in the level of deTyr-tub in MDA-MB-231 cells. We have previously shown this tubulin PTM positively regulates cytoskeletal stiffening in muscle cells as determined by both AFM and indirect measures of passive cell mechanics. [39], [40]. Given our work showing elevated levels of deTyr-tub increase metastatic phenotypes, we used the MCDS to directly assess the impact of elevated deTyr-tub on the cellular mechanics of suspended cancer cells. Indeed, the MCDS identified a significant reduction in cell deformability of the MDA-MB-231 cell line overexpressing VASH1/SVBP (Figure 5B). Taken together, these results further confirm the efficacy of the MCDS, while supporting the concept that cellular mechanical properties may be useful prognostic indicators of cell pathology.

A major strength of the MCDS is its high throughput nature, capturing cells at a rate of hundreds of cells per second. This efficiency highlights time intensive semi-manual analysis methods as major bottlenecks to the full potential of the MCDS. Future work should be done to implement a machine learning algorithm, such as U-net segmentation, to expedite the analysis process [19]; however, in the current work, we intentionally chose to use an ImageJ-based analysis method that is readily understood and simple to implement into almost any lab. Given that manual analysis across several individuals yielded qualitative comparable results that exhibited with detectable variation, (Supplemental Figure 3) all data collected and presented was analyzed by a single individual.

The MCDS was created to function as an accessible and affordable platform for the mechanotyping of single suspended cells on a population scale. We note that system validation was only with cells often used and examined in adherent culture but can also be examined in suspension. Our future works will extend to peripheral blood mononuclear cells (PBMCs) and other cell types that are primarily in suspension. Here, any differences in cell size or dynamic range of cell deformation may necessitate parameter optimization (i.e., flow channel dimensions, fluid viscosity). In this case, the iteration cycle of optimization would be greatly accelerated with the incorporation of a machine learning data analysis approach.

Overall, we have developed an accessible microfluidic cellular deformation system that has been validated and optimized to produce results like those from commercial products. The MCDS provides researchers with a viable method for collecting empirical data regarding mechanical properties of analyzed cells, defining prognostic implications that underlie pathological phenomena, or screening drugs to modulate cytoskeletal dynamics. The MCDS shows promise as a more accessible method to mechanotype single cells at population level that can be easily integrated into any cell culture lab.

## ACKNOWLEDGEMENTS

The authors acknowledge funding from an NIGMS MIRA #R35GM142838 (to KMS) and from the Clark Doctoral Fellowship (to IMS and SB) and NIA P30 AG028747 to the University of Maryland Claude D. Pepper Older Americans Independence Center Scholar and Pilot awards (JU, JS, IMS, KS, and CW). This work was also made possible through the use of the Maryland NanoCenter and its fabrication equipment. The authors also thank Dr. Gregg Duncan’s lab for the generous use of the viscometer. Research reported in this publication was supported by the National Institute of General Medical Sciences of the National Institutes of Health under Award Number R35GM142838. Support was also provided through the National Cancer Institute awards R01-CA124704 and P30-CA134274. The content is solely the responsibility of the authors and does not necessarily represent the official views of the National Institutes of Health.

## Notes

### Competing Interest Statement

The authors have declared no competing interest.

