## Supplemental Figures for "High throughput cell mechanotyping of cell response to cytoskeletal modulations using a microfluidic cell deformation system"

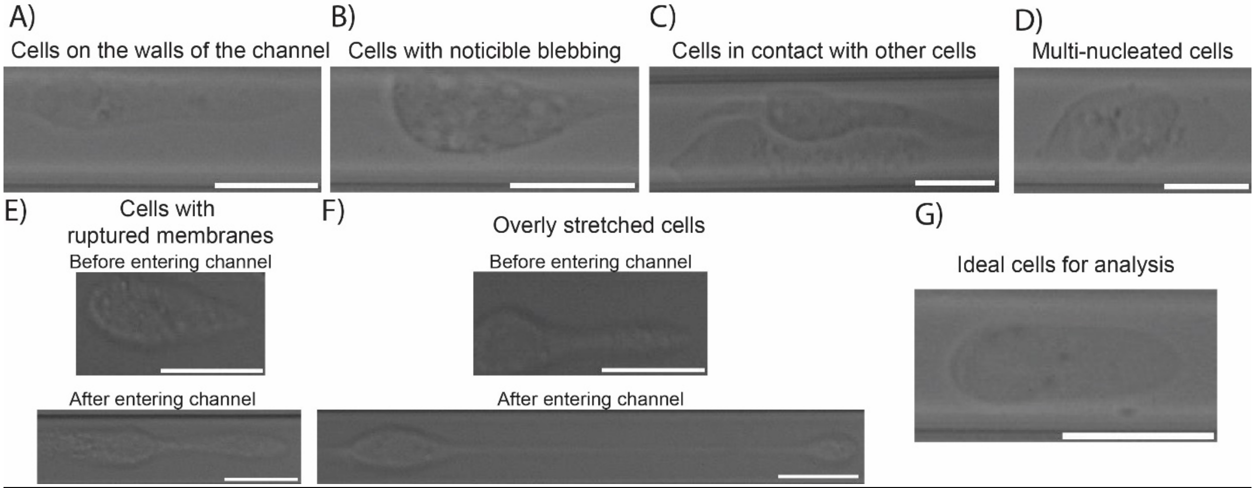


**Supplemental Figure 1: Representative cells manually excluded from analysis.** A) Cells that are dragged along the walls of the deforming region of the MCDS. B) Cells that possess noticeable blebbing as large visible bubble-like structures. C) Cells that are in contact with other cells in any capacity. D) Cells that noticeably have multiple nuclei. In the case of initial trials where cells experienced extensive shear stress, E) cell membranes would visibly rupture as cells traveled through the channel, resulting in a spillage of cellular contents; F) the differences are undefined between stretched cells and ruptured cells, though stretching is seen at the front of cell, while rupturing often occurs at the end. G) A representative image of an "ideal" cell in terms of meeting the criteria needed to be analyzed. Scale bar = 25 µm.

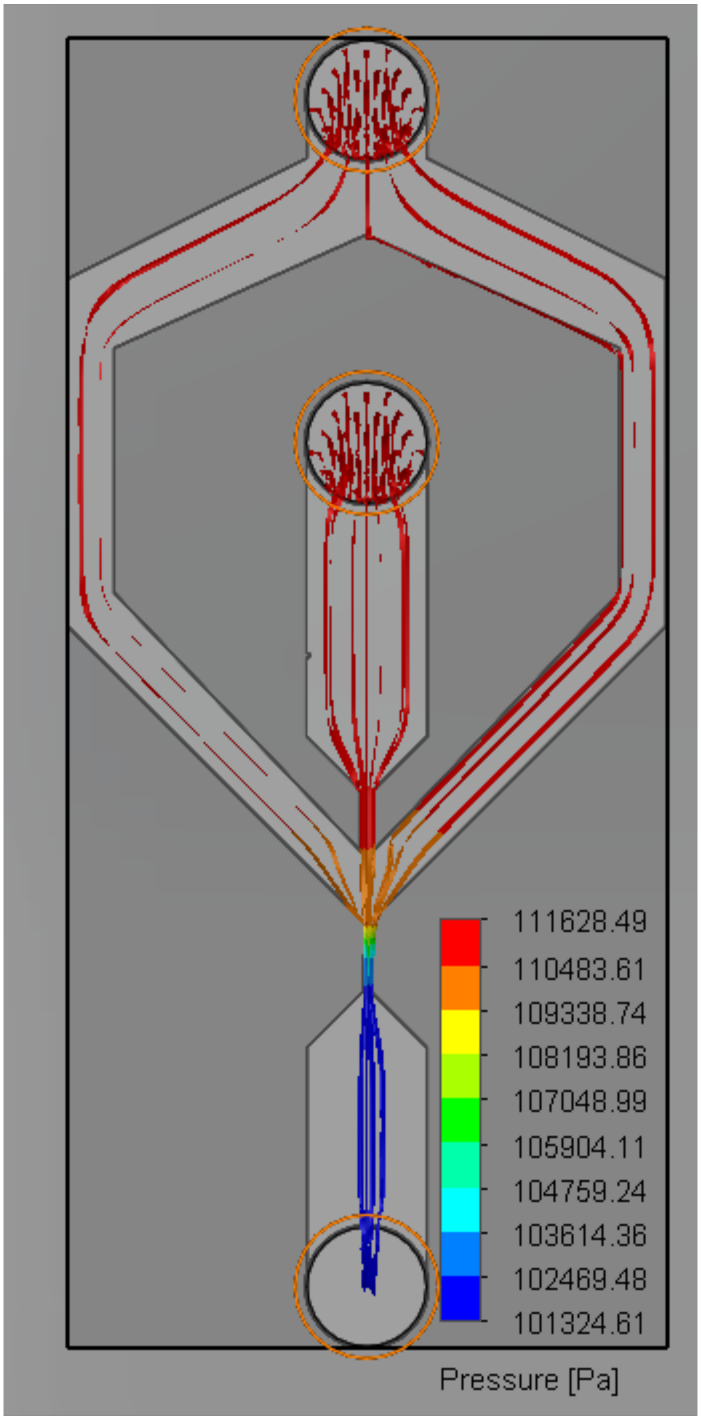


**Supplemental Figure 2:** A SolidWorks FlowView rendition of the pressure at different streams within the SAVS under the optimized flow rates of 300 μL/hour for both the cell suspension and deformation fluid inlets.

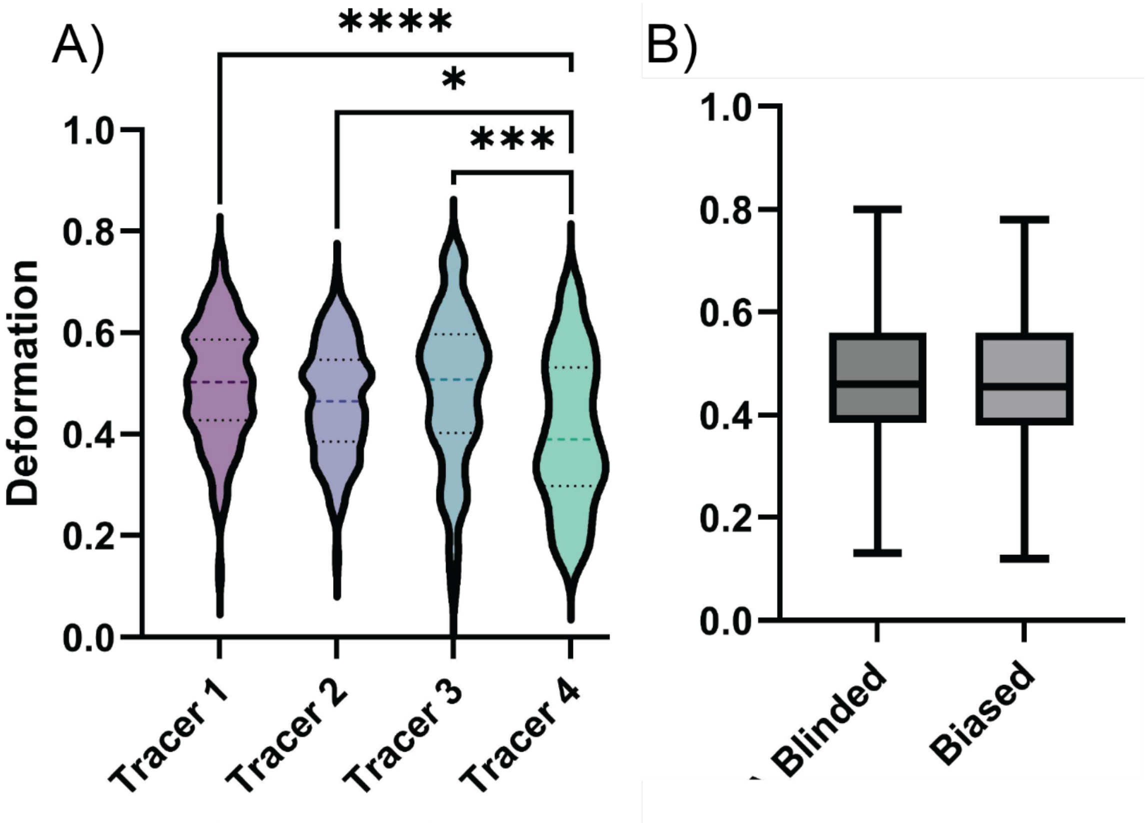


**Supplemental Figure 3: Characterization of blinded vs. unblinded analysis of deforming cells.** A) Five cell videos from a single treatment type were chosen, blinded, and analyzed by five authors. (n=125). B) A treatment was analyzed blinded and then with bias; this produced no significant difference. Violin and box-and-whiskers plots display the min-to-max distribution of data, with a center line at the mean. ****: $p<0.0001$

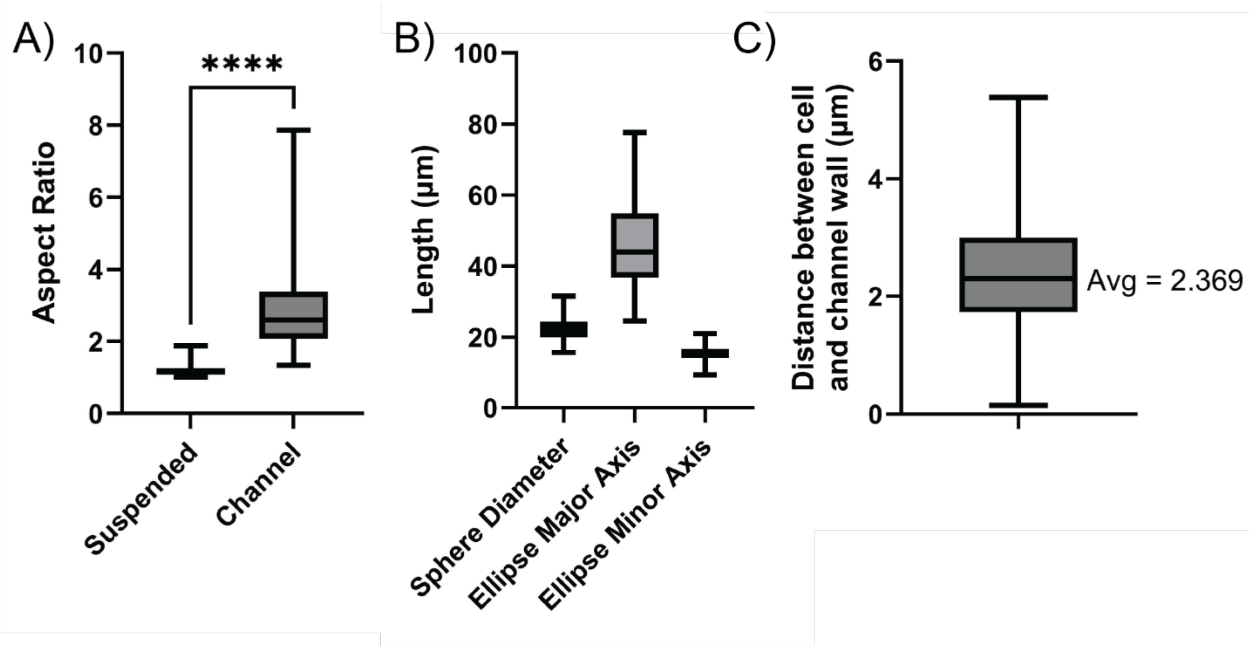


**Supplemental Figure 4: Additional shape descriptors of cells in the deformation region of MCDS.** A) Aspect ratio of cells in the deformation region compared to the same suspended cells. B) Axis from the Fit Ellipse function compared the spheroid diameter of cells in suspension. C) The distance between the edges of cells and the boundary of the channel walls. Box-and-whiskers plots display the min-to-max distribution of data, with a center line at the mean. ****: $p<0.0001$

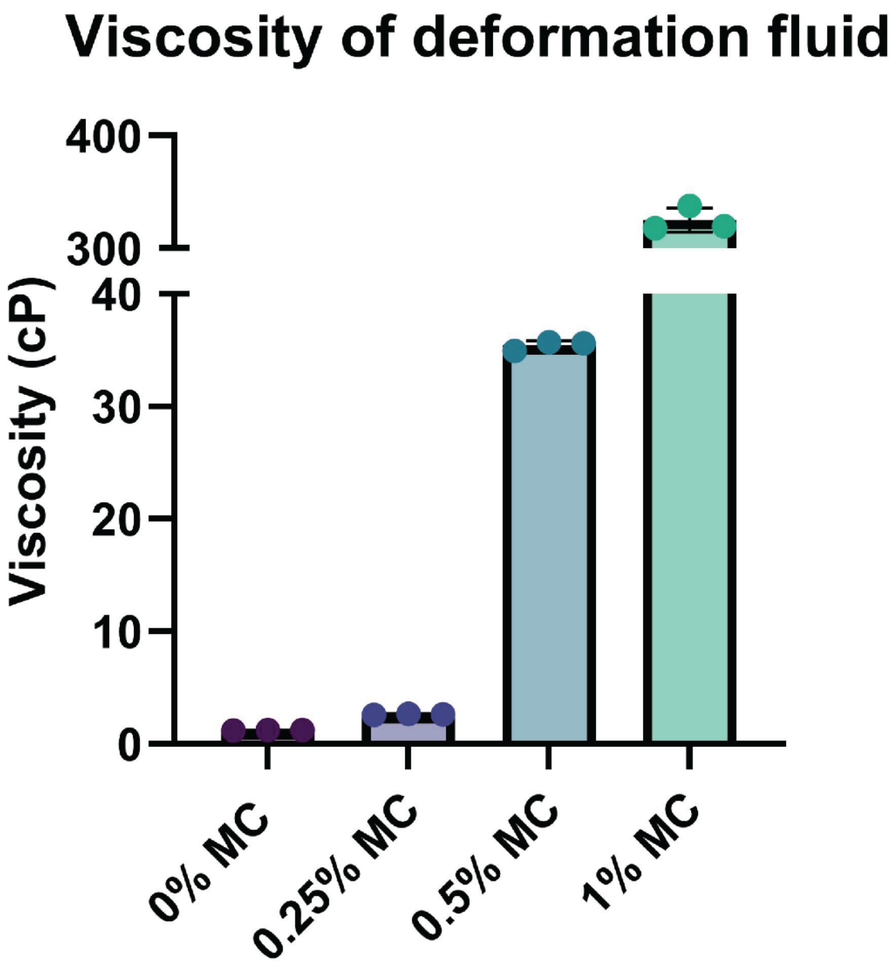


**Supplemental Figure 5:** Viscosity values (cP) of deformation fluids created with varied methylcellulose concentrations. Points represent distinct measurements of deformation fluid viscosity (n=3).

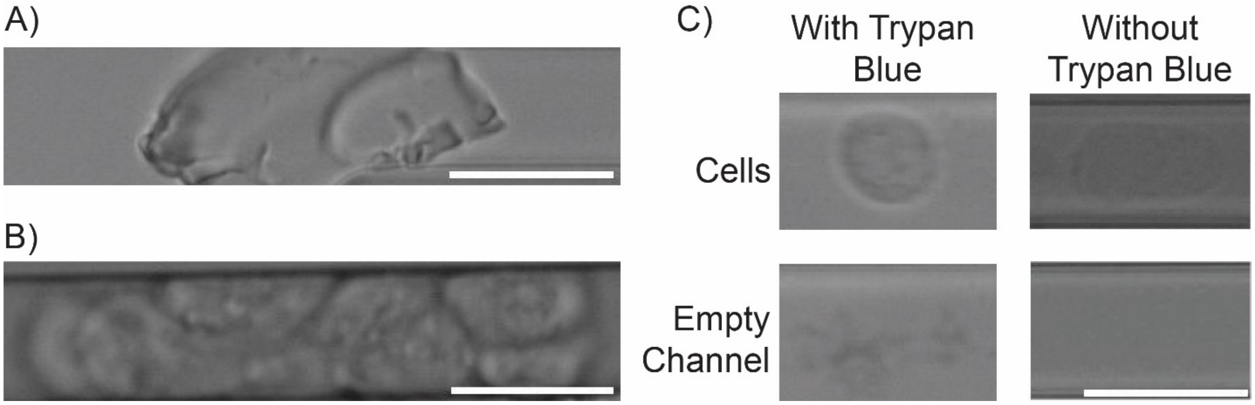


**Supplemental Figure 6: Examples of other optimizations of the MCDS.** A) a representative image of debris trapped at the mouth of the deforming region of the MCDS. Tubing and devices were washed thoroughly in addition to the filtration of deformation fluid prior to running the cell through; this mitigated the blockage of the channel mouth with debris. B) Self adherent cells, such as MCF10As, frequently went through the deforming region in large clusters; these were prevented by straining cells prior to being flown through the MCDS. C) Trypan Blue dye was used in several studies to enhance the contrast of cells. Though it subtly aided in the visualization of cells, it led to an excess of debris as shown in the empty channel panel. Scale bars = 25 µm.
